# ScienceRSI: Recursive Scientific Self-Improvement

**DOI:** 10.64898/2026.10.20.760442

**Authors:** Xun Xu, Jiaxian Yan, Zhicai Wang, Mingyang Xu, Zaixi Zhang

**Author notes:** huggingface.co/Scinetics-ai/ScienceRSI-8b.

## Abstract

Scientific AI can improve cumulatively when capabilities developed in one model become useful learning signals for the next. We introduce **ScienceRSI**, a framework for recursive scientific self-improvement organized around *specialize, verify, transfer, and reuse*. Compact experts develop complementary scientific capabilities; shared evaluation identifies their useful contributions; a target model integrates those contributions and can serve as the starting point for further specialization. This organization connects reasoning, model scaling, and iteration through reusable scientific capabilities. We evaluate its transfer step in **4B** *→* **4B**, *→* **4B***→* **8B**, and **8B** *→***27B** configurations. ScienceRSI ranks first among the compared methods on all five science benchmarks for the 4B and 8B students. Relative to rubric-based reinforcement learning, scientific-reasoning averages improve by 2.0 and 1.9 percentage points, and domain-science averages by 3.3 and 2.1 points, respectively. In a separate cross-generation 27B extension, reported scores improve over Vanilla on all seven benchmarks, with the unweighted mean rising from 87.3% to 94.0%. These results support verified capability transfer across model scales as a practical foundation for ScienceRSI. Accumulating these gains over repeated rounds is the next step toward sustained recursive scientific improvement.

## 1 Introduction

### Making scientific progress reusable

Scientific reasoning combines quantitative calculation, symbolic derivation, mechanistic explanation, and the assessment of evidence. Targeted training can improve each of these capabilities using scientific data and verifiable feedback [1–3]. The broader opportunity is to make those improvements accumulate across model generations. When a specialist learns a useful scientific behavior, its contribution should remain available to the next generalist and to subsequent rounds of specialization.

### The ScienceRSI idea

ScienceRSI organizes scientific model development into a recurring cycle: *specialize capabilities, verify progress, transfer useful knowledge, and reuse the improved model*. Each round produces both a stronger target model and a basis for further learning. This makes scientific capability itself a reusable resource: local advances can be consolidated within a model scale or transferred to a larger model with broader capacity.

### Small experts can contribute to larger models

An expert’s value depends on the scientific requirement it can resolve. A compact model may correctly handle a derivation, mechanism, or evidence check that a larger model’s current answer misses. Recent weak-to-strong learning studies support the feasibility of such transfer [4, 5]. ScienceRSI uses shared scientific evaluation to identify these complementary capabilities before integrating them. The same principle applies to same-scale consolidation and to small-to-large transfer.

### Evidence across three configurations

We study Qwen-family configurations spanning 4B *→*4B consolidation, 4B *→*8B transfer, and 8B*→* 27B transfer. At 4B and 8B, ScienceRSI leads all five science benchmarks among the compared methods while retaining competitive general reasoning. A separate cross-generation 27B comparison reports gains on all seven benchmarks, averaging 6.7 percentage points over Vanilla. Controlled 4B studies further show that selecting useful expert contributions matters: indiscriminate transfer can weaken a model, whereas verified, targeted transfer improves it.

### Contributions

We present a framework for cumulative scientific capability development, evaluate its transfer step at three model scales, and examine the conditions for useful capability integration.

## 2 ScienceRSI Framework

### 2.1 Specialize, verify, transfer, and reuse

ScienceRSI separates capability acquisition from capability integration. Specialists explore different aspects of scientific reasoning, and a target model consolidates their useful progress. The framework has four connected stages:

1. **Specialize**. Develop complementary experts in quantitative reasoning, symbolic reasoning, mechanistic explanation, and evidence assessment.
2. **Verify**. Evaluate expert and target responses against shared scientific requirements to identify where an expert offers a useful contribution.
3. **Transfer**. Integrate those verified contributions into the target model while continuing to optimize its scientific task performance.
4. **Reuse**. Use the improved model as the basis for another round of specialization and integration, potentially at greater model capacity.

Figure 1 places the evaluated model configurations within this framework.

**Figure 1.**
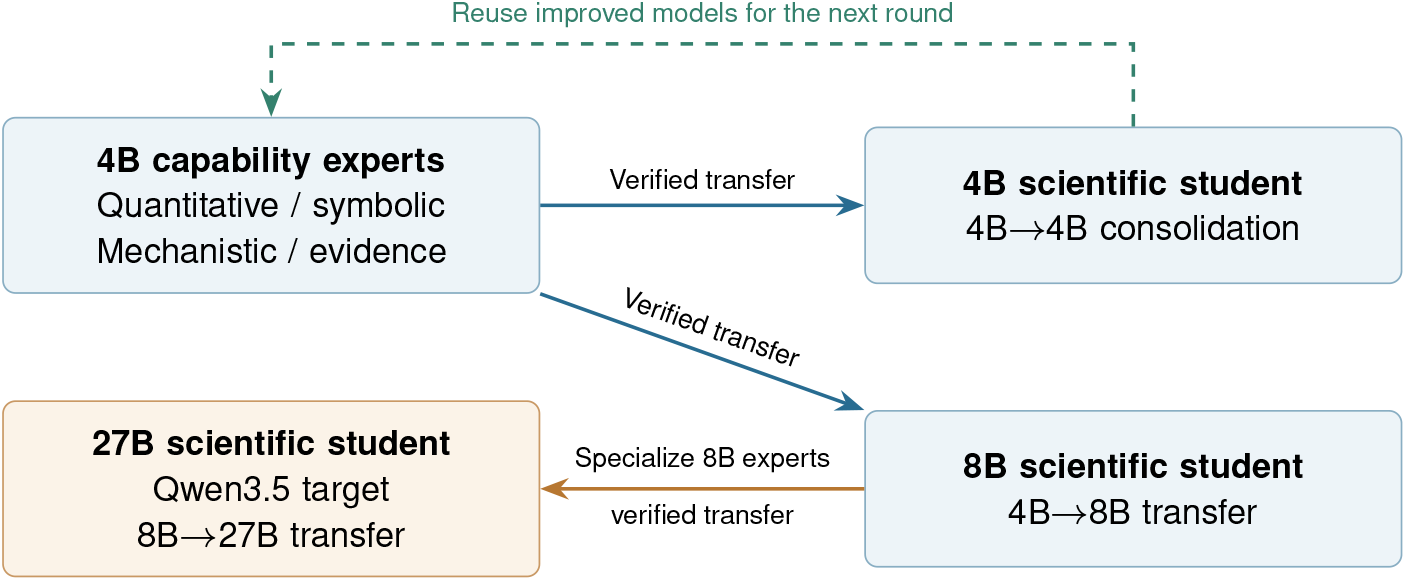
ScienceRSI across model scales. The evaluated settings cover 4B*→* 4B consolidation, 4B *→* 8B transfer, and 8B *→* 27B transfer. Smaller specialists contribute verified capabilities to a target model. The dashed return indicates reuse in the proposed recursive framework; the experiments evaluate separate transfer settings, not successive rounds of one run. The 27B extension also changes model generation.

#### Reasoning, scaling, and iteration

Specialization improves particular reasoning capabilities; transfer makes them accessible to models with broader capacity; reuse creates the opportunity for iteration. Model size can increase between rounds, while same-scale consolidation provides another way to integrate progress. The 4B *→* 4B setting therefore tests capability consolidation, and the 4B*→* 8B and 8B*→* 27B settings test transfer to larger targets.

### 2.2 Scientific feedback connects the stages

Scientific feedback gives the cycle a common basis for deciding what to retain. A response can satisfy some requirements while missing others, and an expert can be useful for one requirement without being uniformly reliable. ScienceRSI compares contributions at this finer level and directs learning toward the parts of a response that can benefit. The target also continues to learn from task rewards, preserving an independent signal of scientific progress. Figure 2 shows the workflow used in the reported experiments.

**Figure 2.**
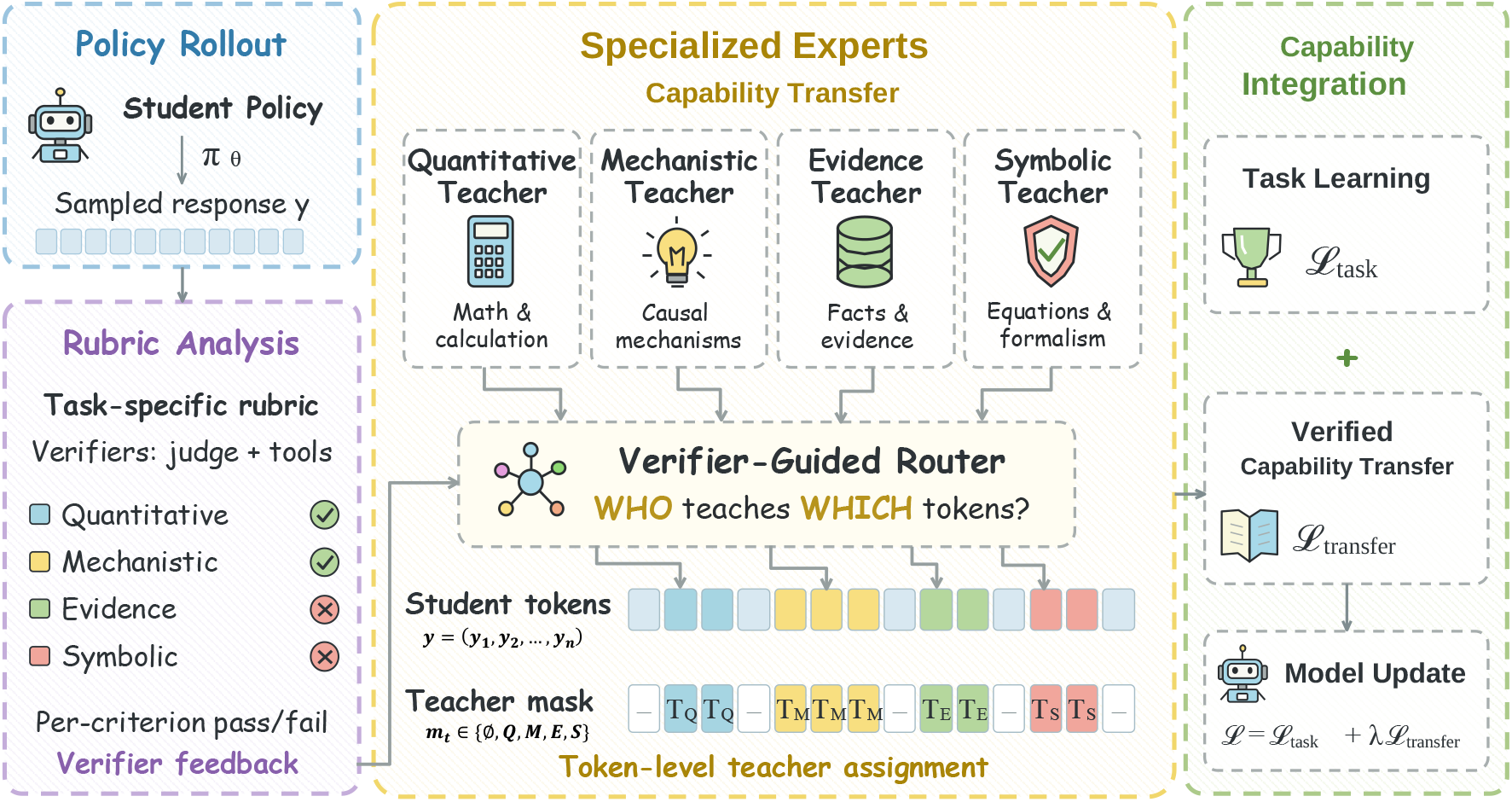
One round of ScienceRSI. Scientific evaluation identifies gaps in a model’s responses, specialized experts provide complementary capabilities, and verified feedback guides capability integration. Task rewards and expert feedback jointly improve the target model. The training formulation is given in Appendix A.

#### From a transfer step to a recurring process

The present experiments evaluate individual transfer settings. In the broader framework, the consolidated model, evaluation criteria, and accumulated feedback support subsequent rounds. This distinction defines a concrete next question: how much useful capability continues to accumulate as specialization and transfer are repeated?

## 3 Evaluation

### Models and scientific capabilities

Qwen3-4B experts supervise Qwen3-4B and Qwen3-8B target models; Qwen3-8B experts supervise a Qwen3.5-27B target [6, 7]. The four expert capabilities are quantitative, symbolic, mechanistic, and evidence-based reasoning. Training combines scientific questions, reference answers, and rubrics from the Dr.SCI data pipeline [3]. Appendices A and B provide the implementation, data construction, and evaluation details.

### Benchmarks and comparison protocol

Seven benchmarks cover scientific reasoning, domain science, and general reasoning (Table 1). At 4B and 8B, the comparison includes the Vanilla checkpoint, rubric-based GRPO, and several capability-transfer baselines under a shared evaluation harness. Distillation-based arms share the data and training budgets; expert-based comparisons use the same expert pool. Training regimes are identified in Table 2: OPSD and MOPD are distillation-only, whereas ScienceRSI combines task rewards with verified transfer. The matched RL-plus-transfer comparison is in Table A4. The 27B study compares the Vanilla checkpoint with the ScienceRSI-trained model as a separate cross-generation extension. Scores are percentages, and group averages weight benchmarks equally.

**Table 1.** Evaluation benchmarks for the 4B/8B comparisons. *n* gives the final held-out subset size; each answer type has one scorer (Section B.2). RaR-Science contains 500 questions after excluding 74 items that overlap training derivatives (Section B.1).

| Group | Benchmark | $n$ | Task form | Scoring |
| --- | --- | --- | --- | --- |
| Scientific reasoning | GPQA-Diamond | 198 | graduate-level multiple-choice questions in biology, physics, and chemistry, written by domain experts to resist answering by search | letter match |
|  | RaR-Science | 500 | held-out subset of open-ended science questions from the Rubrics-as-Rewards collection | item rubric, calibrated judge |
|  | SciBench | 580 | college-level physics, chemistry, and mathematics problems from textbooks with free-form numeric answers | unit-aware numeric match |
| Domain science | ChemBench | 1,391 | chemistry questions across general, organic, inorganic, physical, and analytical subfields; 1,148 multiple choice and 243 numeric | rule-scored |
|  | RaR-Med | 150 | open-ended medical questions from the Rubrics-as-Rewards medicine collection | item rubric, calibrated judge |
| General reasoning | MMLU-Pro | 2,000 | fixed subset of ten-option multiple-choice questions across fourteen disciplines with a reasoning-heavy design | letter match |
|  | MATH500 | 500 | competition mathematics problems with exact symbolic or numeric answers | symbolic equivalence |

**Table 2.** Main results by transfer setting and training regime (scores, %). KD denotes distillation only; RL denotes task-reward learning. OPSD and MOPD here are standalone KD, distinct from GRPO+OPSD and GRPO+MOPD in Table A4. The Qwen3 4B/8B blocks use non-thinking, greedy decoding with a 4,096-token limit. The Qwen3.5-27B block reports a separate cross-generation comparison. **Best** and second-best mark point estimates within each multi-method block, not statistical significance; Δ is the gain over Vanilla.

|  | Method | Regime | Scientific reasoning |  |  | Domain science |  | General reasoning |  |
| --- | --- | --- | --- | --- | --- | --- | --- | --- | --- |
|  |  |  | GPQA D | RaR Sci | Sci Bench | Chem Bench | RaR Med | MMLU Pro | MATH500 |
| 4B→4B | Vanilla | – | 41.4 | 57.9 | 50.8 | 57.8 | 46.9 | 57.2 | 80.9 |
|  | GRPO | RL | 43.9 | 60.6 | 54.5 | 61.2 | 50.7 | 62.5 | **85.1** |
|  | OPSD | KD | 42.1 | 59.6 | 52.1 | 58.9 | 48.5 | 60.2 | 82.8 |
|  | MOPD | KD | 43.8 | 59.0 | 54.0 | 63.1 | 49.4 | 61.6 | 82.4 |
|  | T.A. GRPO | RL+KD | 45.5 | <u>61.2</u> | 51.9 | 62.0 | <u>51.0</u> | **63.8** | 83.6 |
|  | CriPO | RL | <u>45.7</u> | 59.1 | 54.5 | <u>65.5</u> | 50.9 | 62.5 | 84.2 |
|  | Transfer only | KD | 43.4 | 60.7 | <u>55.2</u> | 64.3 | 50.2 | 61.3 | 83.0 |
|  | **ScienceRSI** | RL+KD | **46.3** | **62.4** | **56.2** | **66.4** | **52.2** | <u>63.3</u> | <u>84.6</u> |
| 4B→8B | Vanilla | – | 41.9 | 59.7 | 55.2 | 64.6 | 53.8 | 64.1 | 83.6 |
|  | GRPO | RL | 45.8 | 64.5 | 57.4 | <u>69.3</u> | 57.4 | **67.1** | <u>85.4</u> |
|  | OPSD | KD | 42.2 | 60.6 | 55.9 | 66.5 | 55.4 | 64.9 | 84.2 |
|  | MOPD | KD | 44.1 | <u>66.2</u> | 56.1 | 67.2 | 56.9 | 65.6 | 84.8 |
|  | T.A. GRPO | RL+KD | 45.4 | 65.5 | 56.8 | 68.1 | 57.1 | 66.0 | 83.4 |
|  | CriPO | RL | 45.8 | 64.8 | <u>57.6</u> | 66.7 | <u>57.8</u> | 65.2 | 84.2 |
|  | Transfer only | KD | <u>46.3</u> | 65.9 | 57.2 | 67.0 | 56.2 | 65.8 | 84.5 |
|  | **ScienceRSI** | RL+KD | **47.3** | **67.0** | **59.1** | **70.9** | **60.1** | <u>66.7</u> | **85.7** |
| 8B→27B | Vanilla | – | 85.5 | 89.7 | 86.4 | 87.2 | 82.9 | 86.1 | 93.2 |
|  | **ScienceRSI** | RL+KD | **90.2** | **95.3** | **93.1** | **95.5** | **91.7** | **94.0** | **98.3** |
| | $\Delta$ vs. Vanilla | – | +4.7 | +5.6 | +6.7 | +8.3 | +8.8 | +7.9 | +5.1 |

## 4 Results: Capability Transfer Across Scales

### 4.1 Same-scale consolidation and small-to-large transfer

Table 2 reports all seven benchmark scores for the three configurations. **ScienceRSI leads the five science benchmarks at both 4B and 8B**. Compared with rubric-GRPO, the 4B model gains 2.0 points on scientific reasoning and 3.3 points on domain science. The 8B model gains 1.9 and 2.1 points, respectively. These results show that specialized scientific capabilities can be consolidated at the same scale and transferred to a larger target.

#### General reasoning remains competitive

The general-reasoning average is 0.2 points above GRPO at 4B and within 0.1 points at 8B. ScienceRSI also attains the highest 8B MATH500 point estimate.

### 4.2 Extending transfer to a 27B target

The Qwen3-8B*→* Qwen3.5-27B extension improves on the Vanilla baseline across all seven benchmarks. The unweighted mean rises from 87.3% to 94.0%, an increase of 6.7 percentage points. Gains span 4.7–8.8 points despite the stronger starting checkpoint.

Scientific-reasoning improvements are +4.7 on GPQA-Diamond, +5.6 on RaR-Science, and +6.7 on SciBench. Domain-science gains are +8.3 on ChemBench and +8.8 on RaR-Med. General reasoning also improves, by +7.9 on MMLU-Pro and +5.1 on MATH500. Because both target size and model generation change, this extension is considered separately from the Qwen3 4B/8B comparisons. The 4B experiments provide the controlled component analyses.

### 4.3 What makes scientific capability reusable?

#### Select contributions that resolve real gaps

The 4B analyses show that simply combining experts is insufficient. Dense expert supervision can reduce scientific reasoning below the RL-only baseline, while verified and targeted transfer raises all three group averages (Table A4). Removing verification or assigning experts randomly also weakens the results (Table A5). The quality and relevance of an expert’s contribution are central to effective integration.

#### Apply feedback where it helps

Randomly relocating an equal supervision budget produces the largest loss in the controlled study: 6.3 points on scientific reasoning and 12.1 points on domain science. This supports a practical principle for ScienceRSI: feedback must connect a verified capability to a model’s specific gap. Appendix C reports the detailed ablations, response analyses, and training resources.

#### Combine complementary capabilities

A held-out genetics example illustrates the process (Figure A4). One expert contributes a correct intermediate step, while another contributes the correct final answer. After training, the ScienceRSI model satisfies all seven rubric criteria on that question. The figure illustrates criterion-level routing and the final verifier verdicts. It does not identify which individual training update caused each improvement; aggregate comparisons provide the broader performance evidence.

## 5 Discussion and Outlook

### Toward cumulative scientific capability

ScienceRSI provides an organizing principle for scientific model development: progress made by specialists can be verified, integrated, and reused. The three evaluated configurations study this capability-transfer step in same-scale and small-to-large settings. They do not yet demonstrate compounding gains across recursive rounds. The longer-term opportunity is to make each generation a stronger starting point for the next, so that specialization and scaling reinforce one another over successive rounds.

### Expert headroom can disappear

The 4B thinking-mode pilot (Section A.6) exposes a boundary of reuse: when the current student already matches the available experts on the verifier’s criteria, few contributions are licensed. Repeating the same expert construction and tasks may therefore stall. New data, harder verifiable tasks, more diverse expert training, or stronger feedback are possible ways to restore complementary capabilities; their effectiveness over multiple rounds remains to be tested. Gate acceptance and expert– student gaps should be tracked alongside benchmark scores.

### Scientific feedback defines the scope of progress

Reliable evaluation is central to this process. The current suite covers text-based physics, chemistry, biology, medicine, and mathematics. Extending the cycle to molecular structures, figures, simulations, and experimental observations requires appropriate criteria and feedback for each modality. Such feedback could connect scientific reasoning to richer scientific representations and, eventually, to experimental outcomes.

### Next steps

Repeated-round experiments can test whether gains continue to accumulate, which capabilities transfer reliably, and when new expert specializations are needed. Repeated seeds and paired item-level analyses are also needed: the present 4B/8B results are single evaluations, and small score gaps are not evidence of statistical significance. The next stage is to study sustained improvement per unit of feedback and compute, including the cost of constructing experts, verification, and target training across the complete cycle.

## A Training Implementation

This appendix documents the implementation evaluated in the report. The formulation supports the ScienceRSI transfer step; the main text presents its role in the broader recurring framework.

**Figure A1.**
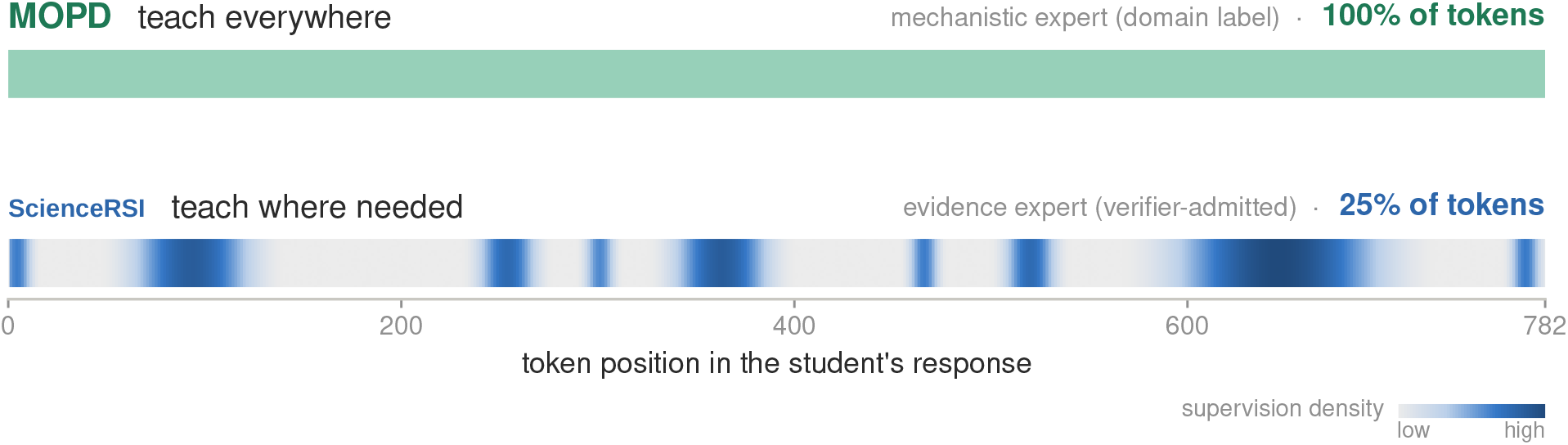
Who teaches which token? Token-level supervision comparison on one 782-token student rollout. MOPD applies dense supervision from a single domain teacher, while ScienceRSI selectively assigns experts and concentrates supervision on tokens where the selected expert disagrees with the student. In this example, the evidence expert supervises only a subset of tokens, with darker spans indicating stronger supervision.

### One rollout through the workflow

Figure A1 shows the outcome of this stage on one 782-token student rollout. Under MOPD, the prompt’s domain label selects the mechanistic expert before the rollout is observed, and that expert supervises all 782 tokens at equal weight. Under ScienceRSI, four intermediate quantities lead to the lower strip. (i) The per-criterion verdicts turn the scalar reward into a set of failed criteria, which says *what* the rollout still lacks. (ii) The capability probes—each expert’s own greedy answer to the prompt—are scored by the same verifiers, which says *which expert* can supply it; here the verifier admits the evidence expert, whereas the domain label points to the mechanistic expert. (iii) The admitted expert’s log-probabilities of the student’s tokens give its disagreement mask, which covers 25% of the response. (iv) The verified weights scale the distillation advantage on the masked tokens. Tokens outside the mask carry no teacher signal and are trained by the task reward alone. Appendix A defines each quantity, and Section C.3 traces a second example down to the answer text.

### A.1 Problem formulation

#### Rubric-based RL yields failed criteria

For each prompt *x*, a rubric defines a set of verifiable criteria 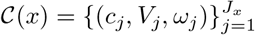, where *c*_*j*_ is a criterion, *V*_*j*_(*x, ·*) *∈* [0, 1] its verifier, and *ω*_*j*_ its signed weight, negative for pitfall checks. Following Rubrics-as-Rewards [2], the rubric is reduced to a scalar reward,

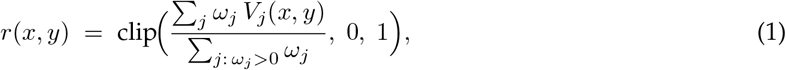

and GRPO [1] optimizes it: a group of *G* rollouts 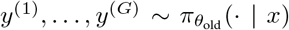 is scored with *r*, the group-normalized reward becomes a shared advantage for every token of a rollout, and the policy takes a clipped step,

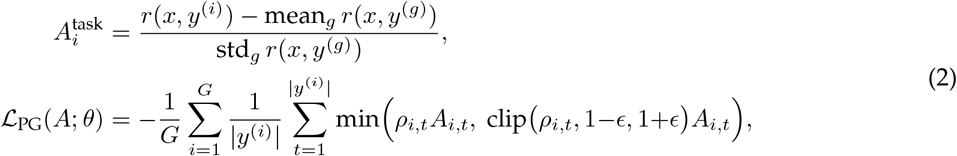

with *ρ*_*i,t*_ the token importance ratio to 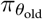 and 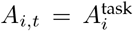 GRPO minimizes ℒ_PG_(*A*^task^; *θ*). The scalar discards structure the rubric provides for free: because *r* is a sum over criteria, every rollout comes with its set of *failed criteria ℱ* (*y*) = {*j* : *ω*_*j*_ *>* 0, *V*_*j*_(*x, y*) = 0}, and the same *V*_*j*_ can score any other answer to *x* on the same criterion.

#### MOPD is fixed routing with dense weighting

On-policy distillation matches the student to a teacher *E* at every generation step along the student’s own rollouts [8, 9]. It yields a token-level distillation signal 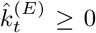, a single-sample estimate of the per-position reverse KL, which enters the same policy-gradient surrogate as the task reward as an advantage 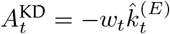, with *w*_*t*_ *≡* 1 in plain OPD [10]. Multi-teacher OPD merges *K* RL-trained specialists {*E*_*k*_*}*into one student; MOPD [11] does so by assigning each prompt the expert of its domain label *d*(*x*) and weighting every token equally:

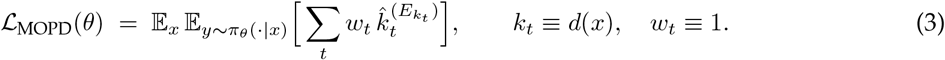

MOPD is therefore a special case of *weighted token-level OPD*, indexed by a teacher assignment *k*_*t*_ and a weight field *w*_*t*_ ∈[0, 1], in which routing is fixed before the rollout is observed and all tokens receive equal weight.

#### From fixed to verified assignment

The rubric says which criteria a rollout fails; the weighted OPD family says who teaches which token and how much. ScienceRSI keeps the objective of Equation 3 and replaces its two fixed choices with verification-guided teacher routing and sparse token weighting: for every failed criterion of every rollout it decides *which expert may teach, which tokens*, and *with what strength*. Only the student’s parameters *θ* are optimized; the experts and the verifiers stay fixed.

### A.2 Who teaches: licensing and routing

#### Licensing

Each criterion and each expert carries a capability axis—quantitative, symbolic, mechanistic, or evidence—written *a*(*j*) and *a*(*E*_*k*_). For a rollout *y* with failed criteria ℱ(*y*) and *j ∈* ℱ (*y*), let *K*(*j*) = {*k* : *a*(*E*_*k*_) = *a*(*j*)} be the candidate experts of its axis. Each candidate produces a *capability probe y*_*k*_ = *E*_*k*_(*x*): a plain greedy answer to the original question, generated once per expert per prompt, cached, and reused across all criteria and all *G* rollouts of the group. The *counterfactual gain* of expert *k* on criterion *j*—the verifier score its own answer receives on a criterion the rollout failed—is

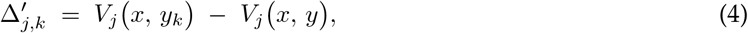

and *k* is *licensed* to teach 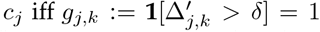. The gate measures, per criterion and per prompt, the condition Li et al. [12] identify for OPD—that the teacher offer a capability the student lacks—instead of assuming it: since *V*_*j*_(*x, y*) = 0 on every failed criterion, the rollout enters the gate only through ℱ (*y*), and one cached verdict *V*_*j*_(*x, y*_*k*_) serves all *G* rollouts of the group. Three exclusions keep the gate honest: pitfall criteria (*ω*_*j*_ *<* 0) never gate; verifier abstentions never gate; and rollouts whose task reward is already high enough receive no teacher term. A criterion with no licensed expert receives no distillation signal at this step. Weak experts are thereby rejected rather than averaged in.

#### Routing

Among licensed candidates the router assigns

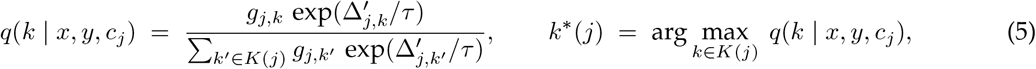

with temperature *τ* =0.5; *q* ≡ 0 when no candidate is licensed. With one expert per axis, *K*(*j*) is a singleton and *q* reduces to the gate; the softmax covers pools with several experts per axis. Licensing and routing are per *criterion*, not per rollout. Let 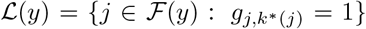 be the licensed criteria of a rollout; its teacher set is (*y*) = {*k*^*\**^(*j*) : *j∈ℒ* (*y*)}. Because criteria of one axis share that axis’s expert, |*T* (*y*) | equals the number of distinct axes among the licensed criteria. For example, a rollout with licensed failures on the quantitative and the mechanistic axis is taught by both experts, each on its own token set.

### A.3 Where to teach: disagreement localization

Each teacher *E*_*k*_, *k ∈ T* (*y*), restricts its distillation to a token mask 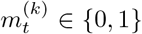 over the student’s own response, computed under its own log-probabilities. Let 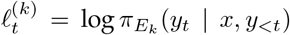 be the teacher’s log-probability of the student’s actual token and 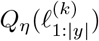 the empirical *η*-quantile of these scores (*η*=0.10); then

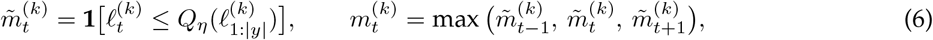

i.e., the positions the teacher “would not have written”, dilated by one token on each side. The undilated seed is rank-based and selects approximately *η* = 10% of the response, with quantile ties and finite-length rounding affecting the exact count. Dilation enlarges that seed. If *N* = |*y*| and 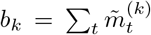, then the single-teacher mask satisfies

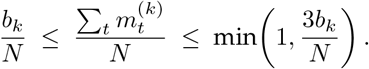

For an approximately 10% seed, coverage is therefore typically between 10% and 30%, depending on adjacency and boundary clipping; the 25% and 27% examples are consistent with this construction. These are conditional, single-teacher mask fractions. Rollouts with no licensed teacher have zero effective coverage; unions of multiple masks and weight-averaged coverage are different statistics. Each mask uses the same teacher forward pass as its distillation target. The gate supplies verifier-based eligibility, not a guarantee that every token in an eligible teacher’s response is scientifically correct.

### A.4 How much: verified-gain weights

For each licensed (criterion, teacher) pair, we assign

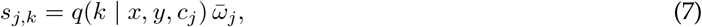

where *q*(*k* | *x, y, c*_*j*_) denotes the verifier-derived teacher eligibility and 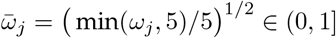 is the criterion weight on the rubric’s five-point scale after square-root compression. The token-level supervision strength and teacher assignment are then

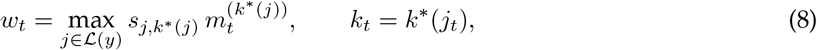

where *j*_*t*_ is the criterion attaining the maximum, with ties resolved in favor of the teacher with larger total verified weight. Thus, each token is assigned to at most one teacher, with its supervision determined by the largest verified weight among the teachers licensed for the corresponding criterion masks; overlapping teacher masks therefore do not produce conflicting gradients. Each masked token is distilled from its selected teacher alone, and tokens outside all masks receive no distillation gradient (*w*_*t*_ = 0).

### A.5 Objective and training step

The teacher term is the weighted OPD of Equation 3, using the per-token teacher *k*_*t*_ and sparse weight *w*_*t*_ from Equation 8. Its per-token signal is

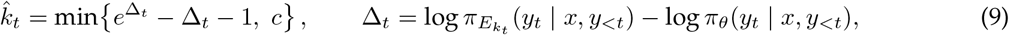

with *c* = 10. Before clamping, and for a fixed teacher and context, the estimator has expectation equal to the reverse Kullback–Leibler divergence 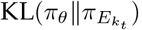; the clamp bounds extreme sampled values. The resulting advantage 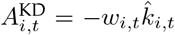 is detached from the gradient, so the auxiliary stream applies a nonpositive penalty to selected sampled tokens, with strength determined by the likelihood-ratio discrepancy. The student is updated with two advantage streams:

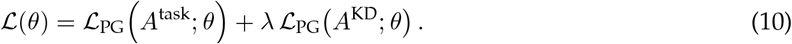

Thus, GRPO reinforces task-level progress, while gated distillation provides targeted teacher pressure where needed. Joint training is important because *A*^KD^ *≤* 0 everywhere:

#### Proposition 1

**(sign and scale of the teacher stream)**

*For every position t*, 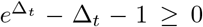, *with equality iff π*_*E*_(*y*_*t*_ | *x, y*_*<t*_) = *π*_*θ*_(*y*_*t*_ | *x, y*_*<t*_), *and* 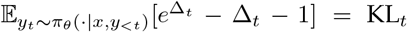. *Hence* 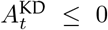 *point-wise*, 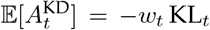 *for a fixed weight before clamping, and the expected teacher pressure on a rollout obeys* 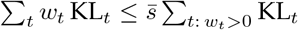 *with* 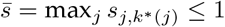, *against* ∑_*t*_ KL_*t*_ *under MOPD*.

The proof is in Appendix E. The sign describes the sampled-token advantage; it does not by itself establish a monotonic change in policy entropy. Distillation can nevertheless overconcentrate the policy. The task-reward stream counterbalances this pressure, while the sparse gate limits it to selected tokens. Algorithm 1 lists one training step; lines marked **+** are the additions to a standard GRPO step.

### A.6 Training strategy and implementation

Each transfer configuration has two stages. Capability experts are trained first, each with RL on its own axis; the target student is then trained with the joint objective of Equation 10 while the experts remain frozen. The student is Qwen3-4B in the 4B *→*4B configuration and Qwen3-8B in the 4B *→*8B configuration, both supervised by Qwen3-4B experts. The 8B *→*27B configuration uses Qwen3-8B experts and a Qwen3.5-27B student (Section 4.2). The implementation details below describe the 4B and 8B comparisons.

#### Capability experts

The four capability experts are rank-64 LoRA adapters on a shared Qwen3-4B base [6], decoded in non-thinking mode. Each is trained with standard GRPO on its axis-filtered data slice: quantitative and symbolic experts use deterministic verifier rewards, while mechanistic and evidence experts use graded judge rewards over imported rubrics. Runs are capped in steps and stopped on sustained validation decline; expert checkpoints are selected by offline re-scoring on the internal dev set. This 4B expert pool serves the 4B and 8B students; the 27B transfer uses the next expert scale, Qwen3-8B.

##### Algorithm 1

Selective capability-transfer training step

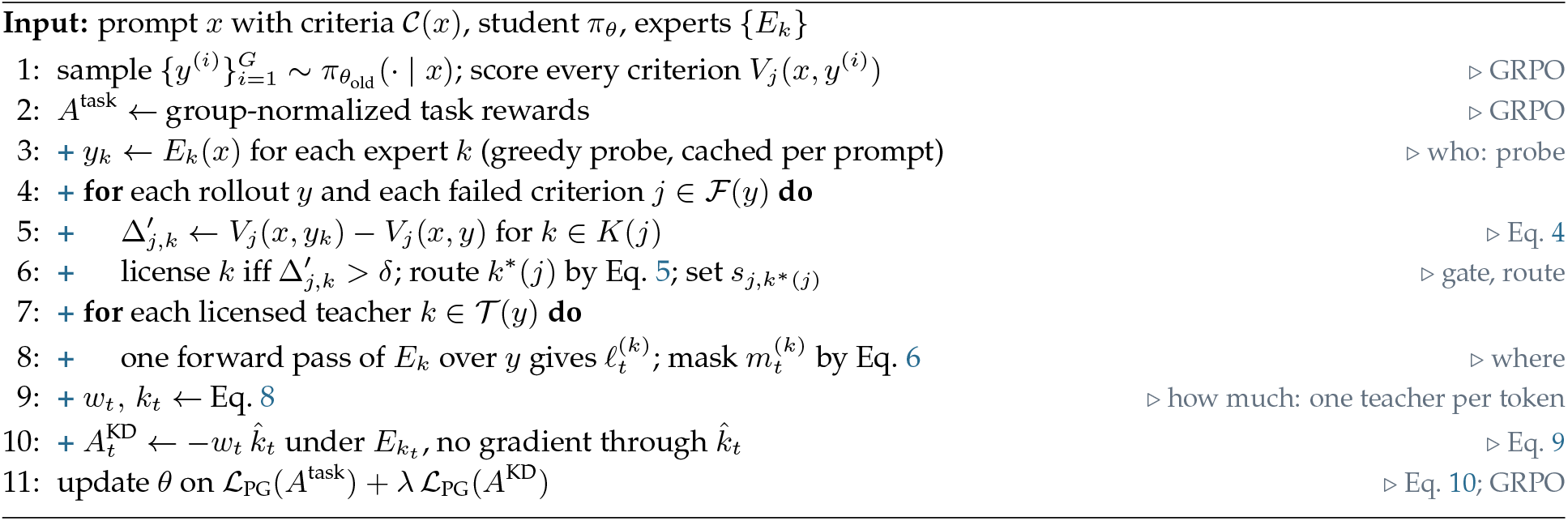

#### Student training

Table A1 lists the shared optimization settings and the distinct teacher–student configurations for 4B*→* 4B and 4B*→* 8B. Baselines share the data and rollout/step budgets but differ in the active objective terms as well as teacher construction (Appendix F). The RL-only floor uses an entropy bonus of 0; the component ablations in Table A5 keep the joint objective and change one transfer component at a time.

**Table A1.**
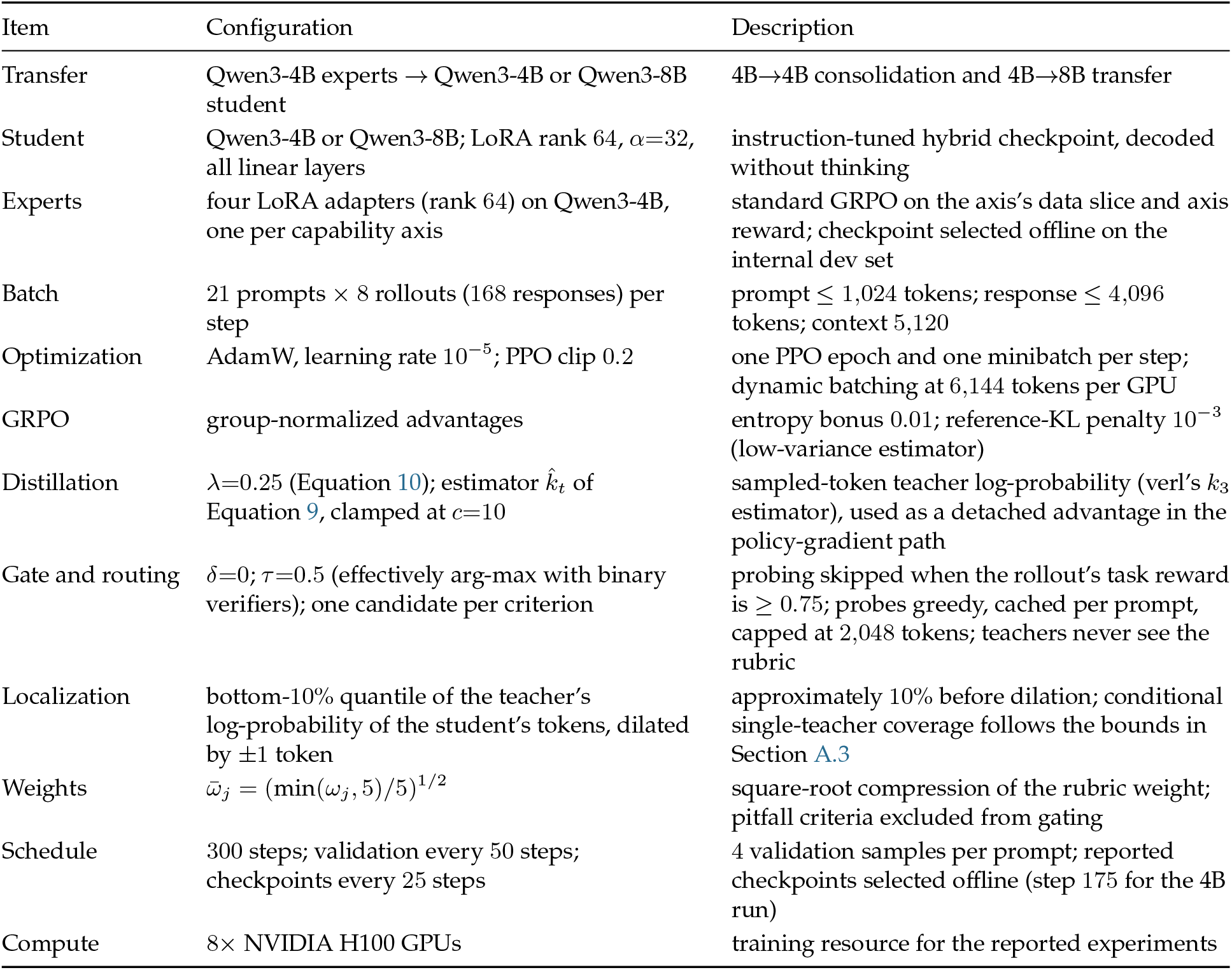
Model and training configuration for 4B*→*4B and 4B*→*8B. Both student sizes use the Qwen3-4B expert pool. Item Configuration Description

| Item | Configuration | Description |
| --- | --- | --- |
| Transfer | Qwen3-4B experts → Qwen3-4B or Qwen3-8B student | 4B→4B consolidation and 4B→8B transfer |
| Student | Qwen3-4B or Qwen3-8B; LoRA rank 64, $\alpha=32$, all linear layers | instruction-tuned hybrid checkpoint, decoded without thinking |
| Experts | four LoRA adapters (rank 64) on Qwen3-4B, one per capability axis | standard GRPO on the axis’s data slice and axis reward; checkpoint selected offline on the internal dev set |
| Batch | 21 prompts × 8 rollouts (168 responses) per step | prompt $\leq 1,024$ tokens; response $\leq 4,096$ tokens; context 5,120 |
| Optimization | AdamW, learning rate $10^{-5}$; PPO clip 0.2 | one PPO epoch and one minibatch per step; dynamic batching at 6,144 tokens per GPU |
| GRPO | group-normalized advantages | entropy bonus 0.01; reference-KL penalty $10^{-3}$ (low-variance estimator) |
| Distillation | $\lambda=0.25$ (Equation 10); estimator $\hat{k}_t$ of Equation 9, clamped at $c=10$ | sampled-token teacher log-probability (verl’s $k_3$ estimator), used as a detached advantage in the policy-gradient path |
| Gate and routing | $\delta=0$; $\tau=0.5$ (effectively arg-max with binary verifiers); one candidate per criterion | probing skipped when the rollout’s task reward is $\geq 0.75$; probes greedy, cached per prompt, capped at 2,048 tokens; teachers never see the rubric |
| Localization | bottom-10% quantile of the teacher’s log-probability of the student’s tokens, dilated by $\pm 1$ token | approximately 10% before dilation; conditional single-teacher coverage follows the bounds in Section A.3 |
| Weights | $\bar{\omega}_j = (\min(\omega_j, 5)/5)^{1/2}$ | square-root compression of the rubric weight; pitfall criteria excluded from gating |
| Schedule | 300 steps; validation every 50 steps; checkpoints every 25 steps | 4 validation samples per prompt; reported checkpoints selected offline (step 175 for the 4B run) |
| Compute | 8× NVIDIA H100 GPUs | training resource for the reported experiments |

#### Starting point

We tested three starts of the same 4B family: a cold-start base with a short SFT stage, the instruction-tuned hybrid model decoded without thinking, and its thinking-mode sibling. In the thinking-mode pilot, the strongest-axis expert’s reward score remained near its initial level (0.585–0.588 over 100 steps at two learning rates). Under the tested settings, expert training produced little verifier-measured headroom over the student, and the gate admitted very few contributions. The non-thinking start improved on the same axis (0.541*→* 0.666 in 200 steps) and is used in the 4B/8B comparisons. This pilot motivates that choice and highlights expert headroom as a constraint on recursive reuse.

#### Checkpoint selection

In-training validation (sampled, mean of four) reordered the final ranking of candidate checkpoints in most arms; every reported checkpoint is the offline winner among the top-*k* in-training candidates under the canonical greedy protocol, selected on the internal dev set before external evaluation.

#### Trainer integration

We build on the distillation stack of *verl* [13] (teacher log-probabilities streamed per token; sampled-token reverse KL with an optional policy-gradient form; multi-teacher routing by a persample key). ScienceRSI adds: (i) a registered distillation loss that multiplies the per-token KL estimate by the per-token weights *w*_*t*_—absent weights fall back to all ones, so every baseline shares one loss implementation; (ii) one hook in the rollout worker, placed after reward computation and before teacher scoring, which parses per-criterion results, runs probes, gate, routing, and localization asynchronously per sample, and attaches the weight vector and, for every token, the log-probability under that token’s teacher to the batch. The hook is fail-open: any internal error zero-fills weights rather than stalling training. Rewards, probe verdicts, and judge calls can overlap rollout generation. This scheduling description does not imply that verification has zero cost.

#### Serving and expert scoring

The expert service shares a base model across its capability adapters. Greedy probes and teacher log-probabilities are computed with frozen experts. For the Qwen3 4B/8B implementation, teacher scoring returns the log-probability of each sampled student token, supplying both the distillation target and the disagreement mask.

**Table A2.** ScienceRSI transfer configurations. All three configurations are evaluated in Table 2.

| Transition | Expert model | Target student | Role |
| --- | --- | --- | --- |
| 4B→4B | Qwen3-4B | Qwen3-4B | Same-scale capability consolidation |
| 4B→8B | Qwen3-4B | Qwen3-8B | Transfer from smaller scientific experts |
| 8B→27B | Qwen3-8B | Qwen3.5-27B | Separate cross-generation extension |

## B Data and Evaluation Details

### B.1 Data construction and splits

#### Training data

Training prompts, reference answers, and open-ended rubrics derive from the Dr.SCI collection [3], via a publicly available reproduction of its pipeline, since the official release was unavailable at the time of writing. Although we use it as one mixture, it aggregates four datasets with different task forms (Table A3). Closed-form items enter the derive-then-verify route described below and open-ended items the rubric-import route. The mixture is non-commercially encumbered: we use it for research only, do not relicense it, and release derived criteria banks under the most restrictive upstream terms. Evaluation sets are public benchmarks used under their own licenses.

**Table A3.** Training data sources. The four datasets are used as one mixture; each expert trains on the slice of its capability axis.

| Source | Content and task form | License |
| --- | --- | --- |
| WebInstruct-Verified [14] | web-sourced questions across physics, chemistry, mathematics, and other disciplines, filtered to those whose short answer (a number, an expression, or a multiple-choice letter) can be checked automatically | Apache-2.0 |
| NaturalReasoning [15] | open-ended reasoning questions extracted from pretraining corpora and paired with model-written reference answers, spanning many domains | CC-BY-NC-4.0 |
| MegaScience [16] | textbook-derived and existing scientific question-answer data across physics, chemistry, biology, medicine, computer science, economics, and mathematics; short-answer and free-form references | CC-BY-NC-SA-4.0 |
| RaR-Science [2] | open-ended science questions paired with expert-guided rubrics | no explicit dataset license; upstream includes non-commercial sources |

#### Criteria bank

Criteria come from two routes. (i) *Verifiable* items (numeric, expression, multiple choice): reference answers are typically bare final values (median ~11 characters), so intermediate criteria cannot be extracted from them. Instead the annotation model solves the problem itself; its derivation is admitted *only if* its final answer passes an endpoint check against the ground truth, after which checkpoint steps become intermediate criteria (restricted to deterministically verifiable types, deduplicated, with target-parsability checks). (ii) *Open-ended* items: the source rubrics are imported verbatim—zero rewriting, audit-friendly— then gated for usability, per-item validity, and axis assignment; style and organization items, which have no capability axis, are dropped; per axis we keep the top-6 items by |*ω*| with the primary axis first. Weights keep their sign; ~16% of stored criteria are penalty (pitfall) items. A row is dropped unless its primary axis has at least one positive-weight criterion, since it would carry no positive learning signal otherwise.

#### Task reward aggregation

The scalar task reward of Equation 1 is computed per capability axis as the signed, weight-normalized sum of criterion scores clipped to [0, 1]: positive-weight criteria contribute their graded verifier score, pitfall criteria subtract |*ω*_*j*_| when triggered, and an axis with no positive-weight criterion scores 0. Verifier abstentions (`unknown`, including criteria left unscored when a sample exhausts its verifier time budget) are excluded from both numerator and normalizer. The final reward multiplies the axis score by a binary format term (a boxed final answer is required), and expert training uses the reward of the expert’s own axis.

#### Verifiers

The registry contains five verifier types:

- `numeric+unit`: parser, unit conversion, and relative tolerance;
- `sympy_equiv`: L^A^TEX-to-symbolic equivalence; a timeout returns `unknown`, which never enters rewards or the gate;
- `sandbox_py`: resource-limited local subprocess re-computation;
- `rule`: constraint and normalized-match checks;
- `llm_judge`: the calibrated judge, for mechanistic and evidence criteria.

Judge calls are cached on (judge version, prompt version, criterion id, answer hash), and the (model, version, prompt version) triple is recorded per verdict. Rollouts must end in a boxed final answer; format failures zero the format term, but open-ended content scoring is not gated on it.

#### Judge calibration

We calibrate the judge by cross-model referee: a stratified sample of (criterion, answer) pairs—two judge axes *×*four subjects *×*positive and penalty items, with answers drawn from the item’s own reference (should pass), a same-subject mismatched reference (fluent hard negative), and a vague template (should fail)—is scored by the production judge under the production protocol and independently by a stronger cross-family referee with full context; disagreements are adjudicated blind by a third model. Outcomes (*n*=578): overall agreement 87.0%; positive-weight items 92.8%; per-axis precision 0.877 (mechanistic) and 0.782 (evidence); vague answers are never credited (0/100), and fluent mismatched references are rejected at 92%. The salient weakness is penalty items: a 39% false-trigger rate on clean answers (the judge over-attributes errors of omission), which is why pitfall criteria are excluded from gating (Section A.2) and carry small |*ω*| in rewards.

#### Evaluation benchmarks and leakage control

We evaluate on seven public benchmarks in three groups (Table 1): scientific reasoning, domain science, and general reasoning. RaR-Science shares an upstream source with our training pool. We removed 74 test items that overlapped training derivatives, leaving a final held-out RaR-Science test set of 500 questions. All reported RaR-Science scores use this post-filter test set (Table 1). The audit also identified a 1,123-item overlap channel between the RaR training split and the training pool. Other benchmarks were checked by exact and prefix matching. Experts and students are selected on an internal dev set held out from the training mixture.

### B.2 Metrics and comparison protocol

#### Metric and aggregation

Every benchmark reports accuracy in percent; higher is better. Multiple-choice answers are scored by letter match, expressions by symbolic equivalence, numeric answers by unit-aware matching with relative tolerance, and rubric items by the calibrated judge against the item’s own rubric. Tables A4 and A5 report *group averages*: the unweighted mean of the benchmark accuracies of a group, rounded to one decimal. Differences quoted in the text are differences of these rounded values.

#### Evaluation harness

The 4B and 8B comparisons use one harness: greedy decoding, non-thinking mode, at most 4,096 new tokens (an 8k budget gave no gain for any arm), identical prompts per benchmark, and one scorer per answer type. Thinking segments, if any, are stripped before scoring; an unclosed thinking segment scores zero. Merged LoRA artifacts are verified by a weights-differ-from-base assertion before evaluation. Each 4B or 8B result is one greedy evaluation of the arm’s offline-selected checkpoint.

#### Baselines and matched budgets

At 4B and 8B, we compare against the starting checkpoint (Vanilla); GRPO with rubric rewards [2], the RL-only floor; OPSD, self-distillation with the rubric as privileged context [17, 18]; MOPD [11]; teacher-anchored GRPO (T.A. GRPO) [19]; and CriPO [20]. All distillation-based baselines share the student, data mixture, rollout and step budgets, and corresponding optimizer settings. The active loss terms differ: standalone OPSD/MOPD and Transfer only have no task-reward stream; ScienceRSI and T.A. GRPO combine RL with teacher supervision; CriPO is rubric-based RL with criterion-specific advantage modification. Sharing experts controls the teacher pool, but comparing standalone MOPD with ScienceRSI changes both task rewards and transfer rules. Table A4 separates these factors by also reporting GRPO+MOPD. Appendix F documents each reproduction.

**Key evaluation detail**

The 4B and 8B comparisons use one harness because the same checkpoint scored several points apart across public harnesses. Rubric benchmarks are judged *serially*: concurrent judging under a per-call time budget deflates rubric scores by up to 0.1–0.2, so every checkpoint was re-scored serially. The RL-only floor runs in its own stable configuration (no entropy bonus) rather than with a copy of the distillation arms’ hyperparameters, which would weaken it (Appendix F).

#### Uncertainty

Each 4B/8B entry is a single evaluation of one selected checkpoint. No across-seed standard deviation or paired confidence interval is available, so rankings are descriptive. On a 198-question benchmark, one question corresponds to approximately 0.51 percentage points; a 1–2-point gap is only about two to four questions. Repeated training runs and paired item-level resampling are needed to assess robustness. Rounded aggregate scores alone are insufficient to recover paired uncertainty.

## C. Additional Transfer Analyses

### C.1 Ablations

Two controlled comparisons on the 4B student ask where the gains come from. Table A4 compares transfer recipes standalone or inside GRPO, with expert-based arms sharing a pool; Table A5 disables one component of the token-weight construction at a time. Figure A2 plots both tables as differences.

**Table A4.** Training-paradigm comparison (group averages, %; 4B student). Expert-based arms share an expert pool; the comparison includes a privileged self-teacher (OPSD), domain-routed experts (MOPD), or criterion-routed and gated experts (ScienceRSI), used standalone or as auxiliary supervision inside GRPO. *Transfer only* is ScienceRSI without task rewards. The standalone MOPD row (52.3/56.3/72.0) matches Table 2; the GRPO+MOPD row (50.1/56.4/73.2) is the distinct model used in the cases. **Best** per column is bold on green.

| Regime | Method | Sci. reasoning ↑ | Domain science ↑ | General reasoning ↑ |
| --- | --- | --- | --- | --- |
| Standalone | OPSD | 51.3 | 53.7 | 71.5 |
|  | MOPD | 52.3 | 56.3 | 72.0 |
|  | Transfer only | 53.1 | 57.3 | 72.2 |
| RL + distillation | GRPO + OPSD | 51.6 | 54.5 | 73.7 |
|  | GRPO + MOPD | 50.1 | 56.4 | 73.2 |
|  | **ScienceRSI** | **55.0** | **59.3** | **74.0** |
| RL only | GRPO | 53.0 | 56.0 | 73.8 |

#### Training paradigm

Selective transfer performs best among the compared recipes in both regimes. Standalone, accuracy rises from OPSD to MOPD to ScienceRSI (scientific reasoning 51.3 *→*52.3 *→*53.1, domain science 53.7*→* 56.3 *→*57.3), and the standalone ScienceRSI variant matches or exceeds the RL-only floor on both science groups without any task reward. Inside GRPO the contrast sharpens: dense distillation drags GRPO *below its own floor* (GRPO+OPSD 51.6 and GRPO+MOPD 50.1 vs. 53.0 on scientific reasoning), whereas gated, localized distillation lifts it by +2.0 / +3.3 / +0.2. These within-regime comparisons support the value of selective transfer beyond merely adding task rewards; the component ablations further separate the transfer choices.

#### Components

Localization matters most: a coverage-matched random mask loses 6.3 / 12.1 / 5.6 points (scientific / domain / general), more than a sequence-level (2.5 / 3.3 / 0.7) or step-level mask (3.5 / 2.8 / 2.8), so distilling the same budget at the wrong positions is worse than distilling everywhere. Who teaches matters as well: random routing costs 5.7 / 4.0 / 3.8 and removing the gate 4.4 / 1.6 / 0.5, both largest on scientific reasoning, and flat weights cost 3.6 / 2.1 / 0.8.

**Table A5.** Component-wise ablation (group averages, %; 4B student). Each variant disables exactly one component of the token-weight construction (Equation 8); data, steps, and *λ* are unchanged. **Best** per column is bold on green.

| Component | Variant | Sci. reasoning ↑ | Domain science ↑ | General reasoning ↑ |
| --- | --- | --- | --- | --- |
|  | **ScienceRSI** | **55.0** | **59.3** | **74.0** |
| Teacher assignment | w/o gate | 50.6 | 57.7 | 73.5 |
|  | Random routing | 49.3 | 55.3 | 70.2 |
| Localization | Sequence-level | 52.5 | 56.0 | 73.3 |
|  | Step-level | 51.5 | 56.5 | 71.2 |
|  | Random localization | 48.7 | 47.2 | 68.4 |
| Weighting | Uniform weighting | 51.4 | 57.2 | 73.2 |

**Figure A2.**
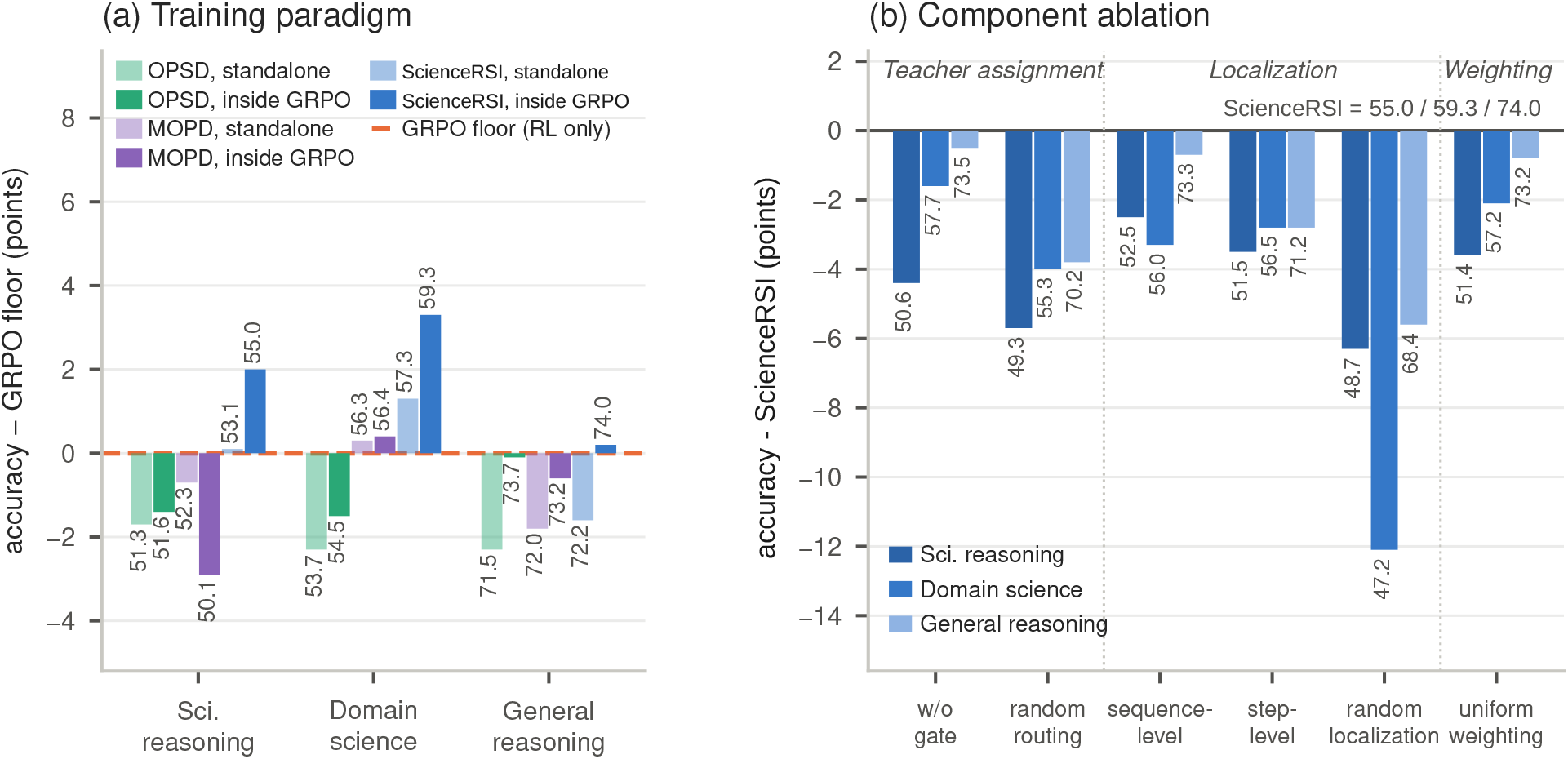
Tables A4 and A5 as differences. (a) Group-average accuracy of each distillation recipe relative to the RL-only GRPO floor (dashed line), standalone (light) and inside GRPO (solid); every bar is labelled with its absolute accuracy from Table A4. (b) Each ablation variant relative to full ScienceRSI, one bar per benchmark group, labelled with the absolute accuracy from Table A5.

In panel (a) of Figure A2, every standalone recipe sits below the GRPO floor on general reasoning and only ScienceRSI reaches it on both science groups; inside GRPO, OPSD and MOPD stay below the floor on every group except MOPD’s +0.4 on domain science, whereas ScienceRSI is above it on all three. In panel (b), the coverage-matched random mask is the largest drop on every group and the only variant that loses more than 10 points anywhere (*−* 12.1 on domain science); the teacher-assignment variants (gate, routing) lose most on scientific reasoning; and flat weights still cost 3.6 points there.

### C.2 Mechanism analysis

**Figure A3.**
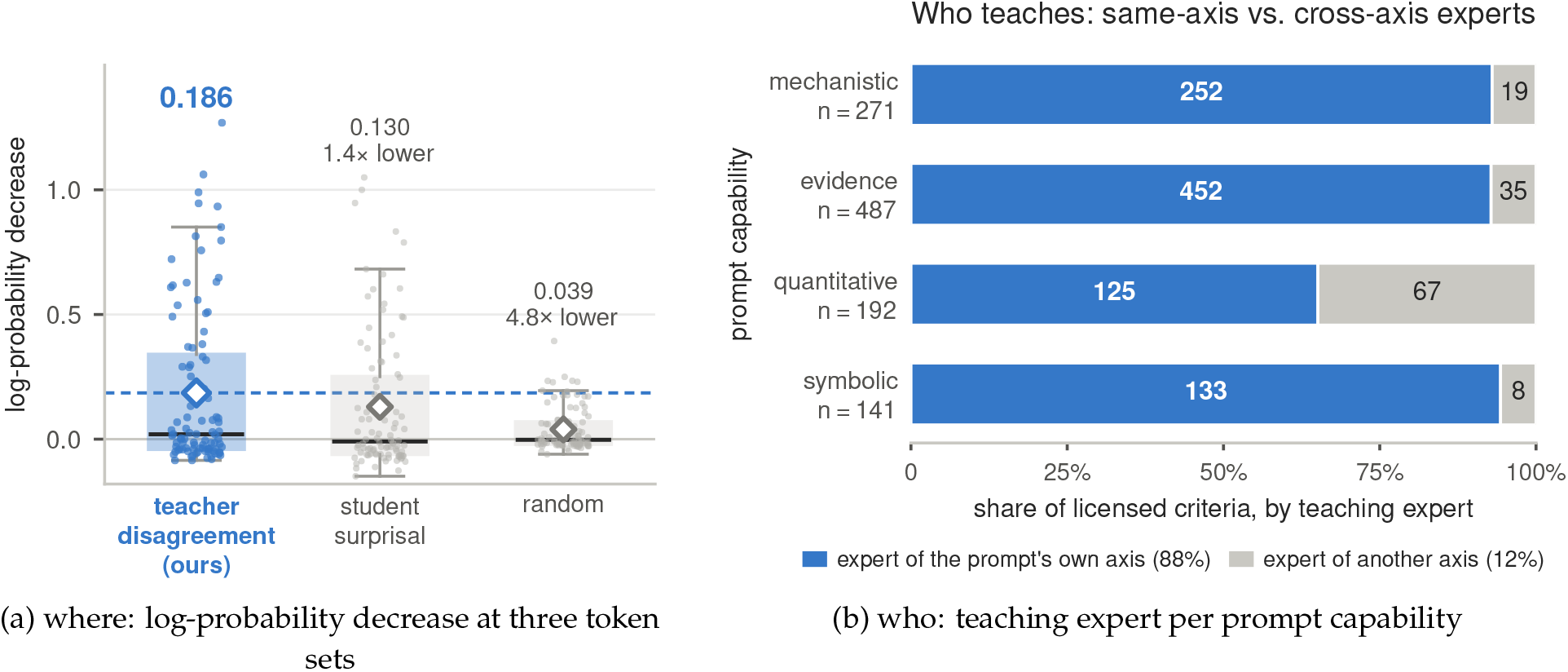
Mechanism analysis. (a) Decrease *D*_*t*_ in the starting model’s log-probability of its own tokens after training, per rollout, at teacher-disagreement, highest-surprisal, and random positions (box: interquartile range, line: median, diamond: mean). (b) Who teaches the licensed criteria: the prompt’s own-axis expert (blue) or another axis’s expert (grey), with counts in the bars.

#### Where: the mask selects corrective tokens

Let *D*_*t*_ = log *p*_start_(*y*_*t*_| *x, y*_*<t*_) *−*log *p*_train_(*y*_*t*_ |*x, y*_*<t*_) be the decrease, after training, in the starting model’s log-probability of its own token. Averaged per rollout over equal-coverage token sets (Figure A3a), *D*_*t*_ is 0.186 at teacher-disagreement positions, 0.130 at the student’s highest-surprisal positions, and 0.039 at random positions. The effect is concentrated: the teacher-marked tokens drop by at least 0.05 in 43% of rollouts, and in 98% of these they drop more than random positions; over all rollouts they drop more than the highest-surprisal positions in 85%. The mask thus concentrates updates on teacher-identified disagreements rather than merely on uncertain tokens [21, 22]. The localization ablations in Table A5 complete the picture: a sequence-level, step-level, or coverage-matched random mask lowers accuracy, and the random mask, which keeps the coverage, lowers it most, so the gain depends on *where* the signal is applied, not on how many tokens receive it.

#### Who: routing is specialization-aligned, not hard-coded

88% of licensed criteria are taught by the expert of the prompt’s own capability (Figure A3b), so routing recovers the specialization that MOPD imposes by label. The remaining 12% cross axes (symbolic criteria of quantitative prompts, mechanistic criteria of evidence prompts) show that the mapping from prompt to teacher is not fixed: a prompt’s failed criteria can call in a second expert.

### C.3 Case study

Figure A4 traces a starting answer with failures on r0, r1, r3, and r4. The mechanistic expert’s separate probe is credited on r1 but rejected on r3–r4; the evidence expert is credited on the final-answer criterion r0. The malformed genotype notation in the middle column is an error in the starting student, so it does not contradict the separately recorded expert verdict.

After the full training run, the verifier credits the ScienceRSI response on all seven criteria. GRPO+MOPD fails r0, r2, r3, and r4: relative to the start, r1 improves but r2 regresses. The latter change is part of the outcome, not evidence of uniformly preserved intermediate reasoning.

#### What the cases do and do not attribute

Licensing describes a particular criterion on a particular training rollout. A post-training response reflects the entire joint objective and many updates, not that one illustrated decision. Improvements on unlicensed r3–r4, or c1 and c5 in Figure A6, may arise from task rewards, shared-parameter generalization, or other training examples; these cases do not distinguish those sources. They are also held-out questions, so diagnostic licensing on the displayed rollout is not evidence that this item was used in a training update. Moreover, a teacher mask is shared across its licensed criteria: rejecting one criterion does not certify that all of that teacher’s erroneous token preferences are excluded. All pass/fail labels are verifier judgments, rather than a separate human validation of every statement.

**Figure A4.**
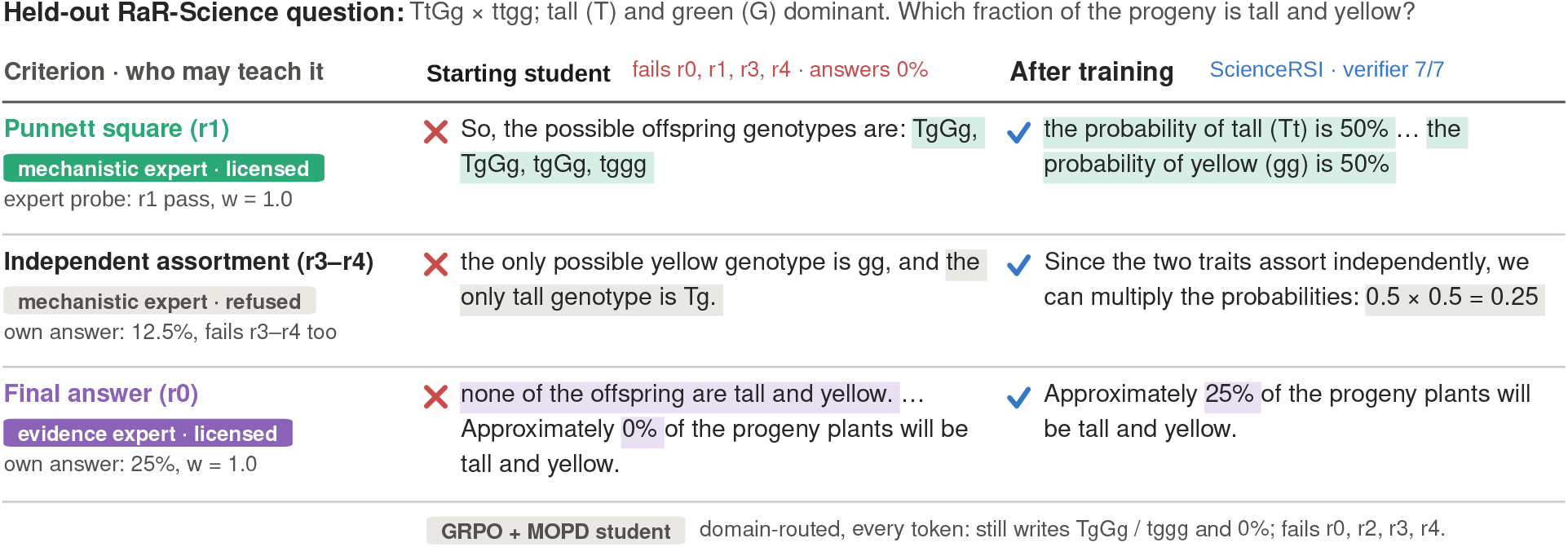
Who teaches which failure. Verbatim student excerpts and verifier judgments on a held-out RaR-Science question. Malformed strings such as TgGg and tggg belong to the incorrect starting/GRPO+MOPD student outputs, not to the mechanistic expert’s probe. The correct offspring classes for *TtGg ×ttgg* are *TtGg, Ttgg, ttGg*, and *ttgg*; tall-and-yellow offspring are *Ttgg* with probability 1/4. The left column summarizes the separate expert probe.

### C.4 Training resources

Training uses 8*×* NVIDIA H100 GPUs.

## D Related Work

### Multi-teacher on-policy distillation

On-policy distillation trains a student on its own rollouts under per-token teacher supervision [8, 9], and its multi-teacher form is now the standard recipe for merging specialists trained by RL. MOPD [11] routes each prompt to the teacher of its domain label and applies uniform sampled-token reverse KL on every token; H-OPD arbitrates among teachers token by token by teacher confidence [23], TU-OPD gates whole examples by reliability scores [24], and CaMOPD addresses recovery–preservation conflicts between teachers [25]. Li et al. [12] show that a teacher helps only where it offers a capability the student lacks; ScienceRSI measures that condition per criterion instead of assuming it.

### Selective and localized distillation

Trust gates decide *when* a teacher may speak: RG-OPD keeps a rollout only when its verifier reward and the teacher–student likelihood gap agree [26], RSTG distills only on prompts whose rollouts all fail [27], SG-OPD gates each token by whether the teacher agrees with the verifier-correct direction [28], SGSD validates the polarity of each skill-conditioned self-teacher against the rollout outcome [29], and TrOPD confines the update to regions where the teacher is reliable [30]. Token selectors decide *where*: decision-critical tokens missed by top-*K* truncation [31], token importance and teachability [21, 22], counterfactual task relevance [32], supervision routed between privileged teacher and student by advantage gap and relative probability [33] or anchored to reference paths [34], and outcome-contrasted patches in Woodpecker [35]. All of these gate or select against a single task-level verdict or an optimization statistic; none checks, per criterion, whether the teacher itself satisfies the criterion it is asked to teach.

### Rubric-based RL for science

Rubric rewards underpin recent scientific RL [2, 3], with criterion-to-verifier routing for robust rewards [36] and automated rubric synthesis [37, 38]. Rubrics also serve as privileged teacher context in self-distillation [17, 18, 39] or replace teacher logits as the scoring signal of on-policy distillation [40], and CriPO decomposes rubric-RL by criterion and self-distills criterion-conditioned revisions with token localization [20]. ScienceRSI instead uses the rubric’s verifiers as a shared test that licenses external experts criterion by criterion.

## E Sign and Scale of the Teacher Stream

The distillation advantage 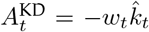 of Equation 9 is non-positive at each sampled position. Policy normalization couples token probabilities, so this sign alone does not determine the entropy of the updated policy. Equation 10 combines the auxiliary stream with task rewards, and the gate and mask restrict its coverage. Proposition 1 states this precisely.

**Proof of Proposition 1**. Non-negativity is *e*^*u*^ *≥* 1 + *u* with equality iff *u* = 0, applied to *u* = Δ_*t*_. For the expectation, 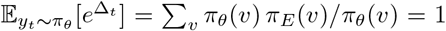 because softmax policies have full support, and 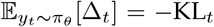 by definition of the reverse KL, so 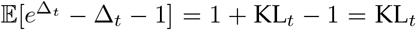. The sign of 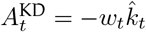 follows from *w*_*t*_ *≥* 0, and the pressure bound from 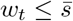 on the mask and *w*_*t*_ = 0 elsewhere; the clamp *c* only lowers 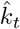 and leaves the sign unchanged. □

## F Baseline Construction

### Fairness protocol

All distillation-based baselines share the student, data mixture, rollout and step budgets, and corresponding optimizer settings with ScienceRSI. Training objectives and teacher construction are explicit comparison factors, not assumed identical. In the 4B/8B main table, MOPD and OPSD are pure distillation stages; Transfer only is the corresponding ScienceRSI ablation. GRPO+MOPD in Table A4 adds task rewards to dense domain-routed distillation and is the model used in Figures A4 and A6. All 4B/8B rows use the same decoding budget, one local harness with serial judging, and offline checkpoint selection (Sections A.6 and B.2).

### GRPO floor

The RL-only floor shares data and budget but disables distillation. The floor uses the *stable-optimal* RL configuration (no entropy bonus), not a copy of our hyperparameters. With the entropy bonus that the distillation arms tolerate, pure GRPO’s entropy diverges because it lacks the mode-seeking KL term that anchors it, which motivates using its stable configuration. Note that the floor is itself a published-method row: its reward is the graded rubric score, i.e., GRPO with Rubrics-as-Rewards [2].

### MOPD [11]

No official code is released; we implement the paper’s policy-gradient form, the sampled-token reverse-KL estimator (their Eq. 4; the paper also reports a top-*k* distillation variant), with the per-token advantage 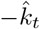 of Equation 9 that every distillation arm in our harness shares (their clipped log-ratio is the *k*_1_ estimator of the same reverse KL), prompt-level deterministic routing by the task’s label (our capability label plays the domain role), uniform token weights, and no verification. Teachers are our four experts—the same pool ScienceRSI uses—so the teacher pool is controlled. The main table reports the pure form (Stage-3: distillation only); comparison with full ScienceRSI also changes the task-reward stream. GRPO+MOPD in Table A4 provides the corresponding joint-training arm.

### OPSD, privileged self-distillation [17, 18]

The teacher is the *student’s own base* conditioned on privileged context: the instance rubric appended to the user message. Teacher log-probabilities are re-indexed onto the student’s unprivileged context position by position; weights are uniform. This isolates “rubric as privileged information” from “verifier-licensed experts”.

### CriPO [20]

We reproduce CriPO: group-level suppressed criteria (their Eq. 4, positive-weight criteria only); a counterfactual self-teacher—the *live* policy served with the criterion removed from context—scored via per-token `prompt_logprobs`; token localization by their Eq. 8 (Δ *>* 0 *∧ p*_*T*_ *< α* max_*v*_ *p*_*T*_, *α*=0.1); and advantage flipping by their Eq. 9 with *τ*_flip_=0.1, applied only where the true advantage is negative. The counterfactual context doubles sequence length, so this arm trains with an enlarged context window.

### Teacher-anchored GRPO [19]

Ramos et al. [19] combine GRPO with on-policy distillation by replacing the reference-policy KL of GRPO with a per-token reverse KL to a stronger teacher, weighted by *β* (their Eq. 6): outcome rewards drive the update while the teacher supplies dense token-level regularization. This baseline does not introduce a separate stronger external teacher, so the row instantiates the recipe with the strongest available privileged teacher—the shared base conditioned on the instance rubric (the OPSD teacher), whose dense term is added to the shared GRPO objective of Table A1 at matched *λ*—and we state this substitution. It isolates “dense privileged fusion” from our gated, localized fusion.

## G Additional Case Studies

One held-out physics question traces the whole mechanism end to end (Figure A6), and a held-out medicine rollout shows the token-level assignment when two experts are licensed (Figure A5). Expert probes and post-training answers are cached greedy outputs; the starting physics rollout is one of four samples drawn at *T* =1 (two already pass every criterion). Verdicts use the training verifier. These held-out examples are diagnostic illustrations of the rule, not evidence of updates on those evaluation items. Questions were selected for clear teacher disagreement and are not a random estimate of error rates.

### Token-level assignment with two licensed teachers (Figure A5)

When a rollout’s failed criteria license two experts, each expert’s disagreement mask is computed under its own log-probabilities (Equation 6) and a token in both masks goes to the larger verified weight (Equation 8). On the medicine rollout of Figure A5 the two masks overlap on 60 tokens, because the experts are adapters on one Qwen3-4B base and find largely the same student tokens unlikely; the assignment leaves the evidence expert 97 tokens and the mechanistic expert 40; their union covers 137/387 *≈* 35.4% of the response. This union across two teachers is distinct from the conditional single-teacher coverage in Section A.3. The second teacher’s exclusive share is therefore modest, and the per-criterion split acts mostly through which criteria license which expert.

**Figure A5.**
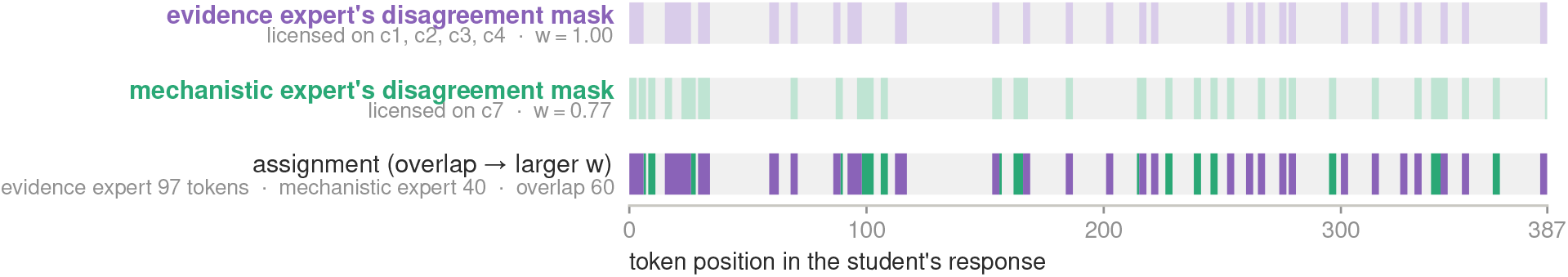
Token-level assignment with two licensed teachers. A held-out medicine rollout (387 tokens) fails the five retained positive criteria, whose source IDs are {c1, c2, c3, c4, c7}. IDs are preserved from the source rubric rather than renumbered: c5/c6 are outside this displayed positive set and do not enter its licensing decisions. The evidence expert is licensed on c1–c4 (*w*=1) and the mechanistic expert on c7 (*w*=0.77). Each expert’s disagreement mask is computed under its own log-probabilities (top two strips); tokens in both masks go to the larger verified weight (bottom), leaving disjoint token sets, each scored by its own teacher.

**Figure A6.**
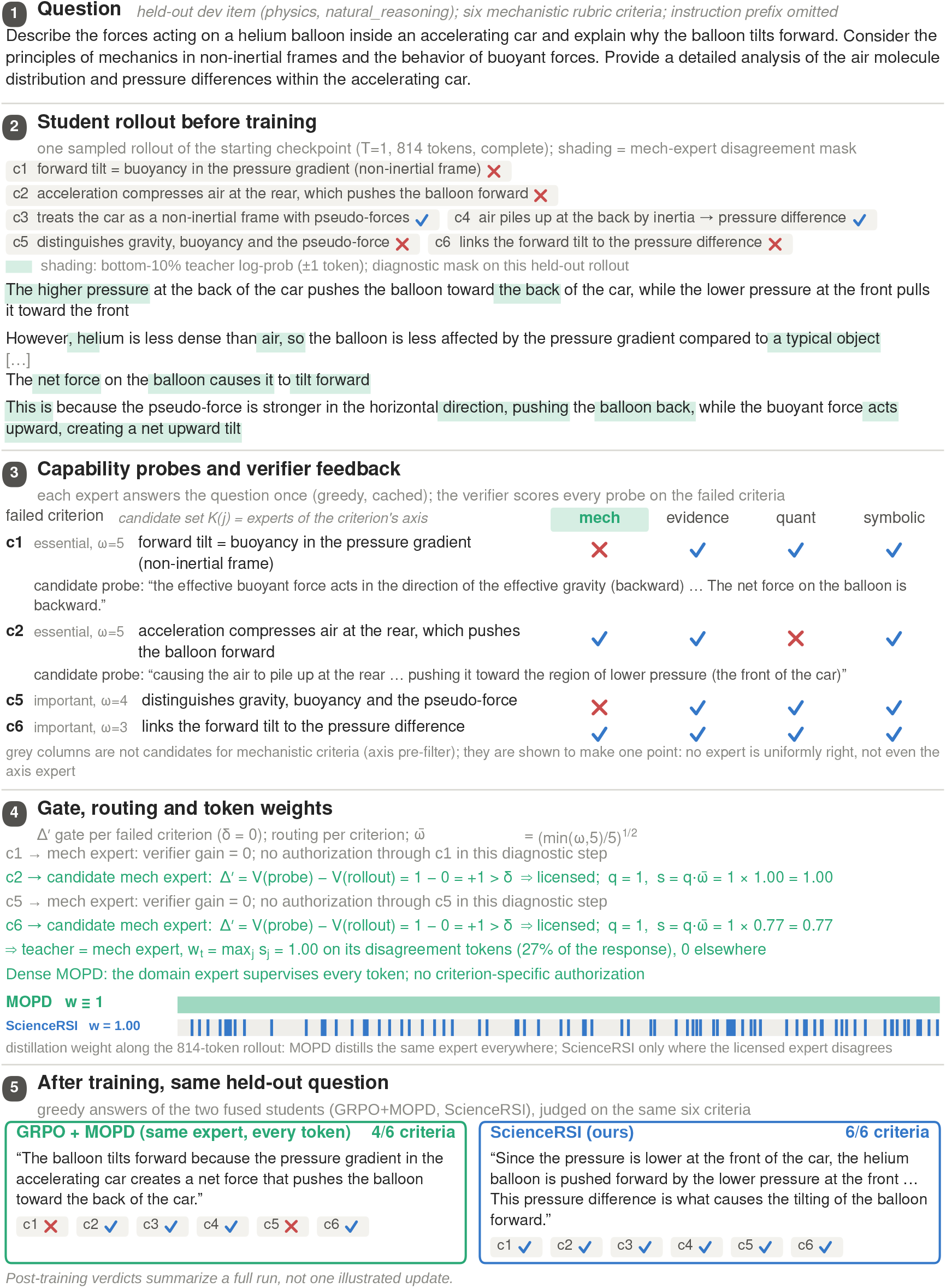
Case study on a held-out physics question (internal dev set). The domain expert is locally wrong: its probe passes the pressure-gradient criteria (c2, c6) but claims a backward net force (c1) and fails c5. ScienceRSI licenses it only where the verifier shows it beats the rollout (c2, c6), refuses it on c1 and c5, and distills the disagreement tokens at the verified weight. After the full run, the verifier credits the ScienceRSI student on all six criteria, including c1/c5, but this case cannot attribute those improvements to a particular reward or transfer update. GRPO+MOPD uses the same expert densely and is credited on four criteria, failing c1/c5. Shading in panel 2 is the diagnostic disagreement mask computed with the training rule; every quotation is verbatim; criterion labels are abridged. The two fused students of panel 5 are the GRPO+MOPD and ScienceRSI rows of Table A4.

## Notes

### Competing Interest Statement

The authors have declared no competing interest.

